# REMAP: An online remote sensing application for land cover classification and monitoring

**DOI:** 10.1101/212464

**Authors:** Nicholas J. Murray, David A. Keith, Daniel Simpson, John H. Wilshire, Richard M. Lucas

**Affiliations:** Centre for Ecosystem Science, School of Biological, Earth and Environmental Science, University of New South Wales, Sydney, Australia; New South Wales Office of Environment and Heritage, Hurstville, New South Wales, Australia

**Author notes:** Corresponding Author:Dr Nicholas Murray, Centre for Ecosystem Science, University of New South Wales, Sydney, Australia 2052, | |.

**Keywords:** Ecosystem monitoring, GIS, Google Earth Engine, Image classification, Landsat Archive, Land cover mapping, Remote sensing, Satellite mapping

## Abstract

1. Recent assessments of progress towards global conservation targets have revealed a paucity of indicators suitable for assessing the changing state of ecosystems. Moreover, land managers and planners are often unable to gain timely access to maps they need to support their routine decision-making. This deficiency is partly due to a lack of suitable data on ecosystem change, driven mostly by the considerable technical expertise needed to make ecosystem maps from remote sensing data.
2. We have developed a free and open-access online remote sensing and environmental modelling application, REMAP (*the remote ecosystem monitoring and assessment pipeline*; https://remap-app.org) that enables volunteers, managers, and scientists with little or no experience in remote sensing to develop high-resolution classified maps of land cover and land use change over time.
3. REMAP utilizes the geospatial data storage and analysis capacity of the Google Earth Engine, and requires only spatially resolved training data that define map classes of interest (e.g., ecosystem types). The training data, which can be uploaded or annotated interactively within REMAP, are used in a random forest classification of up to 13 publicly available predictor datasets to assign all pixels in a focal region to map classes. Predictor datasets available in REMAP represent topographic (e.g. slope, elevation), spectral (Landsat Archive image composites) and climatic variables (precipitation, temperature) that can inform on the distribution of ecosystems and land cover classes.
4. The ability of REMAP to develop and export high-quality classified maps in a very short (<10 minute) time frame represents a considerable advance towards globally accessible and free application of remote sensing technology. By enabling access to data and simplifying remote sensing classifications, REMAP can catalyse the monitoring of land use and change to support environmental conservation, including developing inventories of biodiversity, identifying hotspots of ecosystem diversity, ecosystem-based spatial conservation planning, mapping ecosystem loss at local scales, and supporting environmental education initiatives.

## INTRODUCTION

Maps of land use and land cover change have been a central component of environmental management and conservation planning for decades (Margules & Pressey 2000). Land cover maps enable the depiction of the distribution of ecosystems and land cover types, assessments of biodiversity and identification of areas undergoing loss, fragmentation and degradation (Haddad *et al.* 2015; Potapov *et al.* 2017). As well as supporting spatial conservation planning, including mapping threats to nature, they are often used as surrogates for species distributions. However, existing methods for mapping land cover extent and changes over time are often based on remote sensing and rely on expert implementation and comprehensive knowledge of space borne or airborne sensor data, analytical methods and data uncertainties. This ‘capacity gap’ has been a severe constraint in obtaining information on the status of the world’s natural environment and has hindered environmental conservation programs across a range of spatial scales (Pereira, Brevik & Trevisani 2018; Murray *et al.* in press).

Recent advances in geospatial data access, storage and analysis have vastly improved our ability to utilize satellite sensor data archives in studies of land cover and land cover change (e.g. Lewis *et al.* 2016; Gorelick *et al.* 2017). Moderate (< 30 m) resolution remote sensing analyses are now possible at the global extent and have enabled the development of complex remote sensing analyses (Gong *et al.* 2013; Hansen *et al.* 2013; Pekel *et al.* 2016). At the same time, increases in satellite revisit frequencies, reductions in the time between data acquisition and delivery to users, and increasing access to data archives have led to the development of near real-time alert systems that can rapidly identify land cover loss and change in areas where no ground observations can be obtained. These systems mainly focus on automatic detection and analysis of land cover change for groups of related biomes (e.g. forests) and have vastly improved the ability of non-specialists, environmental managers and policy makers to access and use remote sensing data (Asner *et al.* 2009; Hansen *et al.* 2016; Lucas & Mitchell 2017).

In this manuscript, we present a new online geospatial application that enables volunteers, managers, students and scientists with little or no experience in remote sensing to develop classified maps of land cover at Landsat spatial resolutions. The *Remote Sensing Monitoring and Assessment Pipeline* (REMAP) utilizes the geospatial data storage and analysis capacity of the Google Earth Engine (GEE; https://earthengine.google.com), a cloud-based analysis platform, to allow users to interactively develop machine learning classifications of land cover within an area of interest anywhere in the world for which there is sufficient archival Landsat data. The REMAP application additionally allows monitoring and analysis of land cover change by enabling users to map ecosystem distributions at two points in time (i.e. 2003 and 2017), quantify area change of each map class, and report the standard distribution size metrics used by the International Union for the Conservation of Nature (IUCN) Red List of Ecosystems (Keith *et al.* 2013).

REMAP was developed to complement a range of other applications that support the conservation of biodiversity, including GeoCAT (Bachman *et al.* 2011), Global Forest Watch (www.globalforestwatch.org), the Map of Life (www.mol.org) and R packages such as ‘redlistr’ (Lee & Murray 2017) and ‘rCat’ (Moat & Bachman 2017). Potential uses of REMAP include mapping the distributions of ecosystem types (Murray *et al.* in press), developing land cover maps for protected areas (Lucas *et al.* 2015), assessing the performance of protected areas over multi-decadal time frames (Green *et al.* 2013; Murray & Fuller 2015), and identifying areas where degradation of ecosystems has occurred (Bhagwat *et al.* 2017). REMAP was also developed to support the global effort to assess the status of all ecosystem types on earth under the IUCN Red List of Ecosystems criteria (Keith *et al.* 2015; Rodríguez *et al.* 2015) and can contribute to monitoring progress towards addressing the 2020 Convention on Biological Diversity Aichi Targets (CBD 2014). We describe here the rationale for design, methodological considerations and analytical framework of REMAP, and demonstrate its utility and limitations with four case studies (see *Case Studies*).

### REMAP: REMOTE ECOSYSTEM MONITORING & ASSESSMENT PIPELINE

REMAP (https://remap-app.org) is a free and open-source web application that classifies land cover according to user-supplied training data and a set of globally available remote sensing datasets as predictor variables (Figure 1). We followed six design principles to develop REMAP:

**Figure 1.**
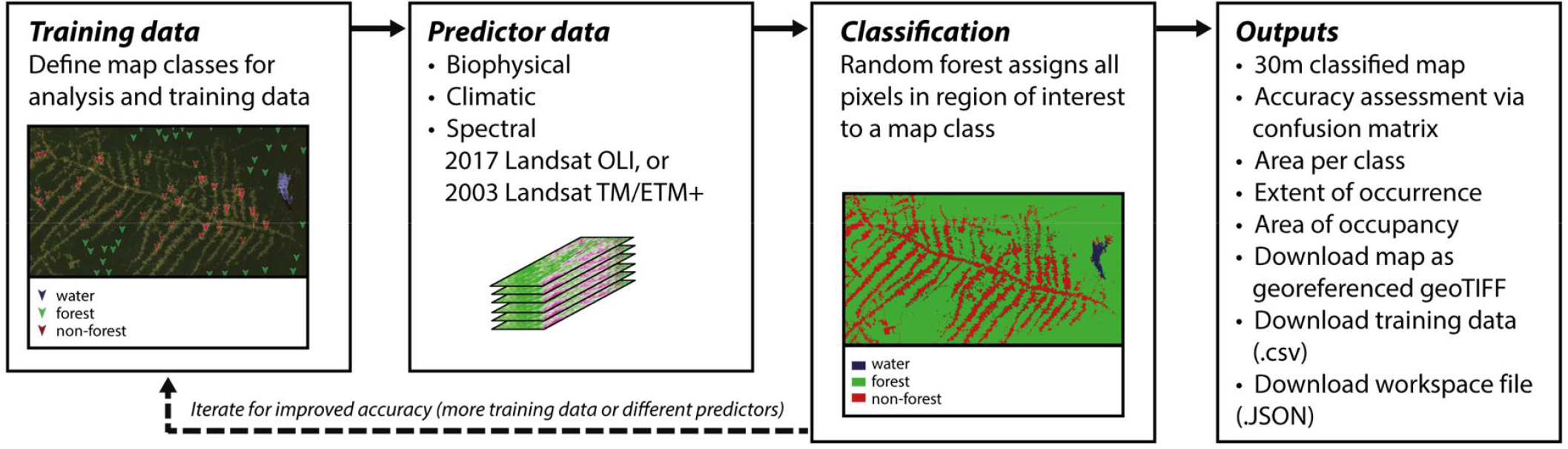
Simplified process chart of REMAP: the remote ecosystem assessment and monitoring pipeline. REMAP requires spatially resolved training data, and estimates class membership of all pixels in a region of interest using global remote sensing predictor layers and the random forests classification algorithm. To facilitate observations of land cover change, classifications in REMAP can be implemented on Landsat data obtained in the year 2003 or data obtained in the year 2017.

1. *Provide the ability to develop high quality maps from remote sensing data in a short time frame and without the need for high performance computers*. Maps can be developed in REMAP within a few minutes and, because REMAP completes classifications online by accessing the GEE, the only prerequisites are an internet connection and web browser.
2. *Reduce the need to download, pre-process and process remote sensing data for use in environmental mapping*. The system offers access to 13 publicly available geospatial predictors that represent spectral, topographic and climatic variables that may influence the distribution of different land cover types. Default predictors were selected to enable the development of high quality maps of the widest range of land cover types possible, and users are provided with options to explore different combinations of predictors in the production of their classified map.
3. *Simplify implementation of machine learning classification approaches*. REMAP conducts its classifications using the random forest algorithm (Breiman 2001) with a single execute button. This approach allows users to implement a widely used machine learning method known to achieve high classification accuracy from large amounts of potentially correlated predictor variables (Rodriguez-Galiano *et al.* 2012).
4. *Permit the production of maps for at least two time periods to enable the quantification of any detectable spatial change*. REMAP can be used to measure the impacts of, for example, deforestation (Hansen & Loveland 2012), coastal reclamation (Murray *et al.* 2014), and many other land cover changes that can be reliably observed with Landsat sensors.
5. *Enable estimation of standard spatial metrics used for assessing the status of ecosystems*. Metrics that are useful for environmental conservation, including area, change in area, extent of occurrence (EOO) and area of occupancy (AOO) can be calculated by users to assess ecosystem change and contribute to global efforts to assess the status of ecosystems.
6. *Implement free and open access software design principles*. Source code for REMAP is available and we will maintain open access to the application (see *Data Accessibility*).

### DATA

The 13 publically available gridded datasets that were selected for inclusion in REMAP (Table 1) met the requirement of (i) full global extent, (ii) free availability with sufficient open access to be included in the GEE public data archive, and (iii) sufficiently high spatial resolution to permit identification of ecosystem distributions and common land cover classes. The final set of predictors includes spectral variables and derived indices from archival Landsat sensor data for two time periods, climate data (mean annual rainfall and mean annual temperature; Hijmans *et al.* 2005) and topographic data (derived from Shuttle Radar Topography Mission data).

**Table 1.**
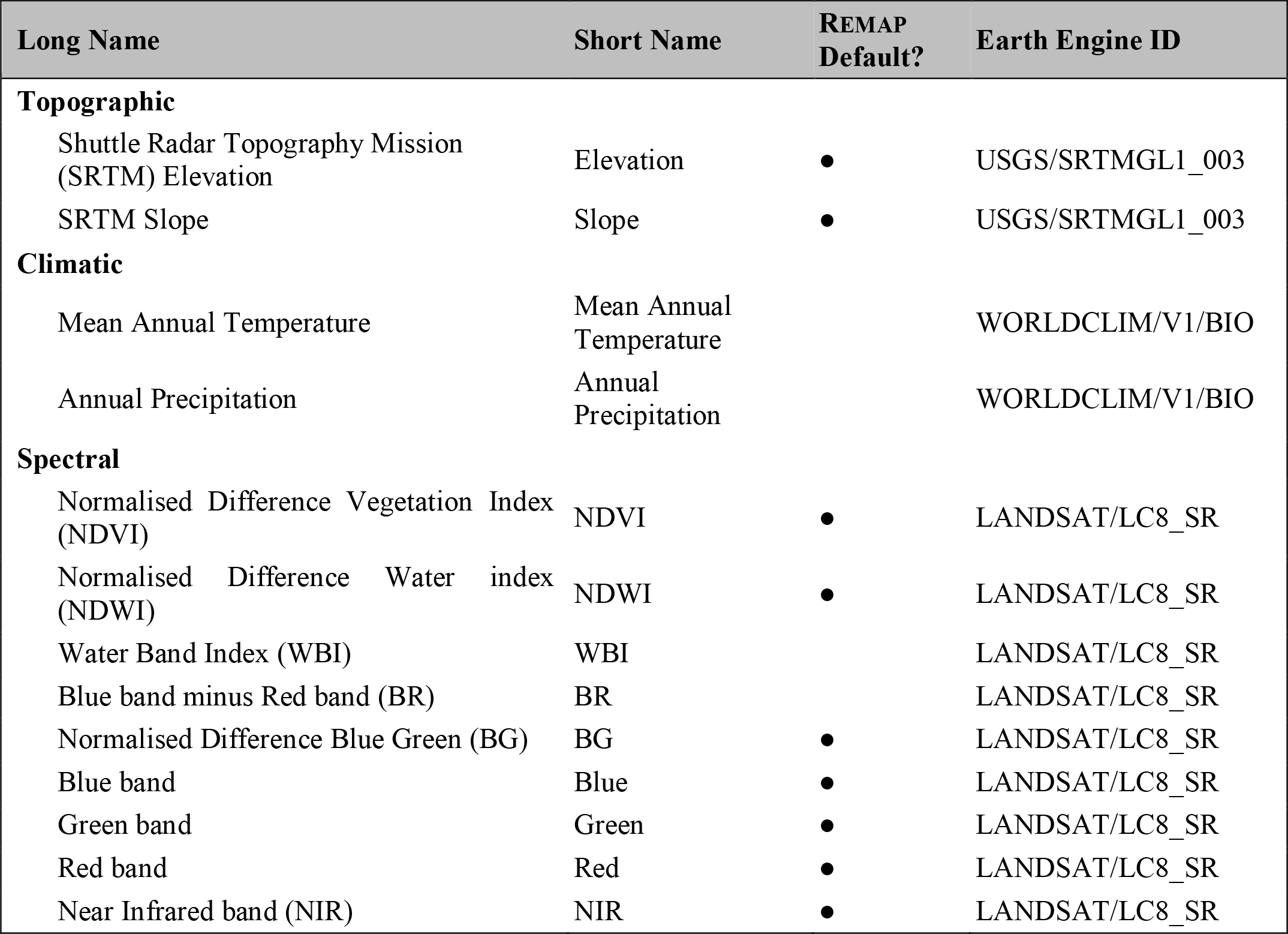
List of predictor layers available for use in land cover classifications using REMAP. Short name refers to the name given to each layer in the REMAP user interface. REMAP default indicates whether the predictor is used in a default classification. Raw data for all predictors used in REMAP are available for download from the Google Earth Engine.

To obtain the required global coverage of cloud-free Landsat sensor data for two periods, referred to here as historical (1999-2003) and ‘current’ (2014-2017), we developed two global Landsat image composites. We produced image stacks of all Landsat scenes for each period (N_1999-2003_ = 340,658 images; N_2014-2017_ = 375,674 images) and applied the GEE implementation of the FMASK cloud masking algorithm (Gorelick *et al.* 2017). From these, the median pixel of Landsat Enhanced Thematic Mapper (ETM+; bands 2-5) bands 2−5 (visible blue to shortwave infrared) and Operational Land Imager (OLI; bands 1-4) was used to generate the two 4-band global image composites. From these composites, Normalized Differenced Vegetation Index (Pettorelli 2013), Normalized Difference Water Index (McFeeters 1996) and several other index layers were generated for use as spectral predictors (Table 1). The provision of spectral data for two time periods facilitates the estimation of change in land cover extent, which is important for monitoring of the impact of threatening processes such as deforestation (Hansen *et al.* 2013), fragmentation (Haddad *et al.* 2015), coastal reclamation (Murray *et al.* 2014), aquaculture (Thomas *et al.* 2017) and water extraction (Tao *et al.* 2015).

### USER INPUT

Users of REMAP generally follow a 7 step procedure to map, assess and monitor ecosystem types or land cover classes (Table 2). Initially, users are required to define their region of interest interactively (focus region) or to upload a vector file (.kml). This enables REMAP to clip input data to a region of interest and limit the extent of the classification. The maximum size of the region of interest is presently 100,000 km^2^ due to limitations applied to users of the GEE (Gorelick *et al.* 2017). Future versions of REMAP may increase this size limit, although for larger regions or more complex map classifications we recommend users directly utilise the GEE (https://earthengine.google.com).

**Table 2.**
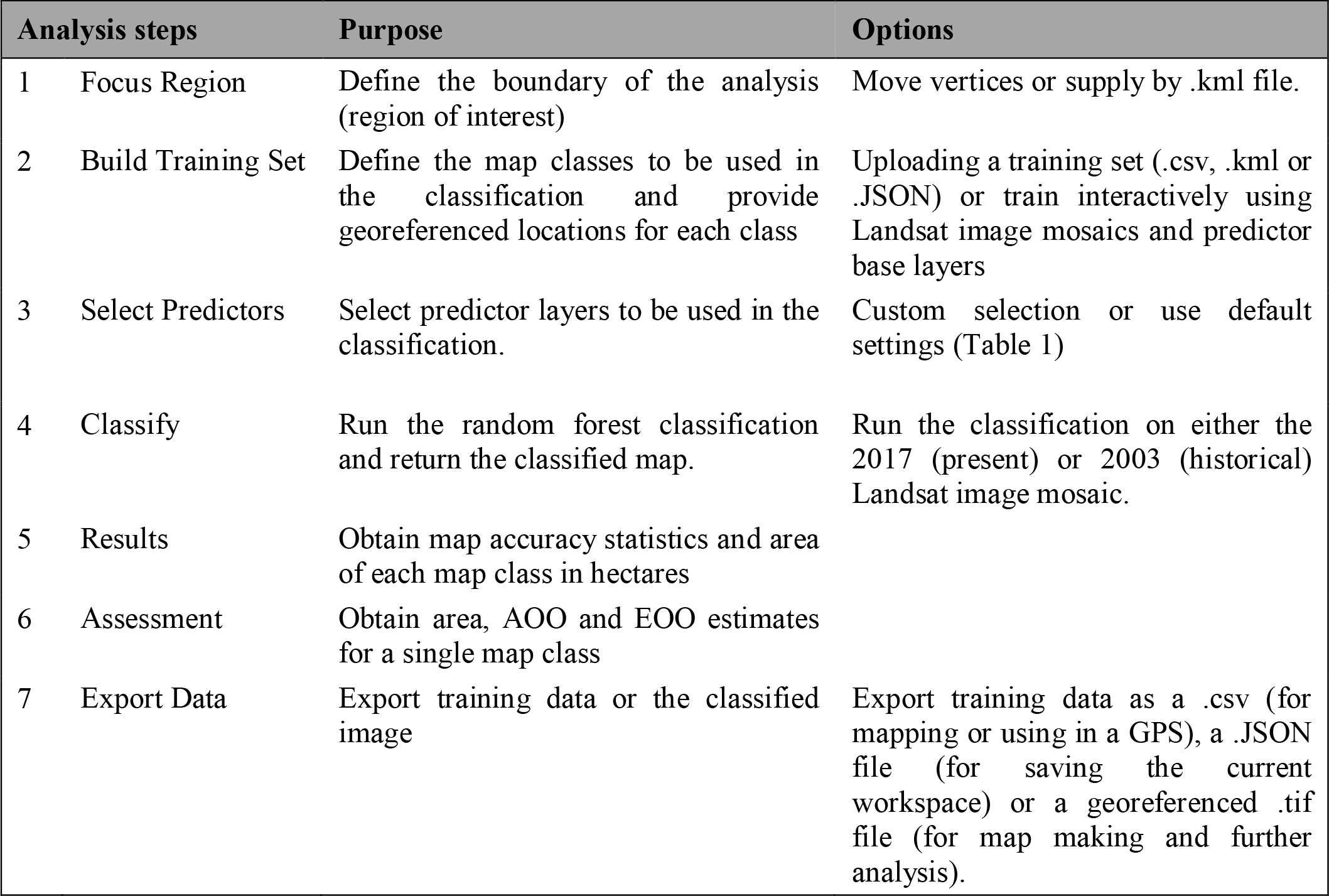
Descriptions of major analysis steps required to develop classified maps in REMAP. Analysis step refers to button in the sidebar of the REMAP user interface.

Spatially resolved training data that define map classes of interest, which can include ecosystem types, land cover classes, areas of change (e.g. deforestation) or anthropogenic areas (e.g. urban areas) are used to assign a class membership to all pixels within a focal region. If developing land cover maps, we recommend that users adopt land cover classification taxonomies that are internationally recognized and confirm to International Organisation for Standards (ISO) such as the Food and Agricultural Organisation’s (FAO) Land Cover Classification System (LCCS). Training data can be provided interactively by adding training points via the user interface with reference to the predictor layers or by uploading data which identify the location of observation points and their class membership (.csv file). These may be sourced from field observations, external data archives, expert opinion, literature or existing maps. In general, classifications with larger numbers of training points will achieve higher class accuracies and we recommend users supply a minimum of 50 points per class to develop an initial map.

### CLASSIFICATION APPROACH

REMAP uses a random forest classifier to assign pixels to user-defined map classes (Breiman 2001). With sufficient training data that are representative of the classes of interest, REMAP implements the classification on the predictor data and returns a classified image to the browser window. In many cases, use of the default predictors (Table 2) will yield classification accuracies that are acceptable to the user. To allow users to assess classification accuracy, REMAP returns a confusion matrix that compares classification results with a random subset of points held-out of the training dataset. Users can tune their classifications to maximize accuracies, either overall or for the class(es) of interest, (ideally to >85%; Congalton & Green 2008) by providing more training data for the classifier or by selecting a custom set of predictors (Table 2).

### ECOSYSTEM MONITORING AND ASSESSMENT

Once a classified map of acceptable accuracy has been produced, REMAP can conduct the spatial analyses required to assess Criteria A (change in distribution size) and B (range size) of the IUCN Red List of Ecosystems (Keith *et al.* 2013; Bland *et al.* 2017). To assess Criterion A, REMAP computes the area of each class by summing the number of pixels in each class. Criterion A requires assessors to estimate change in area over time, which can be achieved by repeating the workflow for the second time period. To account for potential changes in land cover between the two time periods, users should develop a new training set or modify the existing set to ensure accurate representation of land cover in the second time period. Once area estimates are completed for two time periods, assessors can follow the IUCN Red List of Ecosystems guidelines to estimate area change manually (Bland *et al.* 2017) or with the recently developed ‘redlistr’ R package (Lee & Murray 2017). To assess criterion B of the IUCN Red List of Ecosystems, REMAP applies a minimum convex polygon to a class of interest and reports its area, representing the Extent of Occurrence (EOO) of the map class. Finally, the Area of Occupancy (AOO) of a map class is calculated by applying a 10×10 km grid and counting the number of grid cells occupied by the map class (Bland *et al.* 2017; Murray *et al.* 2017).

To support further analyses of the classified map data, users can export each classified map as a georeferenced raster file (.tif). Furthermore, training data can be exported as a .csv file with fields ‘latitude’, ‘longitude’ and ‘class’ suitable for import into a GPS unit or GIS software. Training data can also be saved as a JSON file, which is analogous to ‘save workspace’ functions in other software. This allows users to return to their analysis at a later time by uploading the JSON file (see Appendix 1 for examples).

### CASE STUDIES

Classifications of remote sensing data enable the measurement and monitoring of an enormous range of environmentally relevant variables. To demonstrate the use of REMAP, we developed case studies for (i) mapping a single ecosystem type (e.g. Murray *et al.* 2012; Nascimento *et al.* 2013), (ii) generating a comprehensive land cover map for a region of interest (e.g., Malatesta *et al.* 2013; Connette *et al.* 2016), and (iii) quantifying land cover change between two periods (e.g., Sexton *et al.* 2013; Olofsson *et al.* 2016; Thomas *et al.* 2017). All training data (.csv) and REMAP workspace files (.JSON) used to reproduce these case studies are available in supplementary material (Appendix 1) and can be used in association with tutorials available on the REMAP website (https://remap-app.org/tutorial).

1. *Mapping single land cover types or ecosystem types*. Mapping the distribution and change of mangrove ecosystems has been an important focus of ecosystem monitoring programs for decades due to their provision of ecosystem services (Mumby *et al.* 2004; Spalding *et al.* 2014) and susceptibility to a wide range of threats (Cavanaugh *et al.* 2014; Asbridge *et al.* 2016; Duke *et al.* 2017). In this case study, we developed a simple classification of mangroves and non-mangrove from a set of 150 training points for a small focal region (8301 ha) in the Gulf of Carpentaria, Australia (Figure 2). Against random subsets of training data, the resubstitution accuracy reported by REMAP was 99.2%. Furthermore, a random allocation of 389 points over the focal region indicated a 93.3% agreement with the 2000 global mangrove map data produced for the year 2000 (Giri *et al.* 2011).
2. *Comprehensive classification of land cover for a focal region*. Land cover maps, which represent all land types in a region, is a common aim of remote sensing programs (Lucas & Mitchell 2017). We used REMAP to develop a land cover map with classes *semi-deciduous vine forest*, *eucalypt woodland* and *human settlement* for a focal region in the dry tropics of Northern Australia (Figure 3; Figure S1). A comparison with ecosystem maps produced by the state-wide regional ecosystem mapping program, which develops regulatory land cover maps through manual interpretation of aerial photography and Landsat TM and SPOT satellite imagery, indicated good agreement between the two mapping methods (Figure 3; Neldner *et al.* 2017; Queensland Department of Natural Resources and Mines 2017). We provide a second land cover example that covers a larger area with more land-cover classes in the Supplementary Material (Cheduba Island, Myanmar, Figure S2).
3. *Quantifying land cover change*. To demonstrate capacity to detect changes in land and water, REMAP was applied to the two Landsat composite images available (2003) and OLS (2017) data acquired over Dubai, United Arab Emirates. The resulting maps provide quantitative information on the extent of marine ecosystem loss as a result of large-scale coastal reclamation projects (Figure 4). REMAP’s use for change mapping is also demonstrated with a deforestation example at Roraima, Brazil (Figure 1, Figure S3, Appendix A).

**Figure 2.**
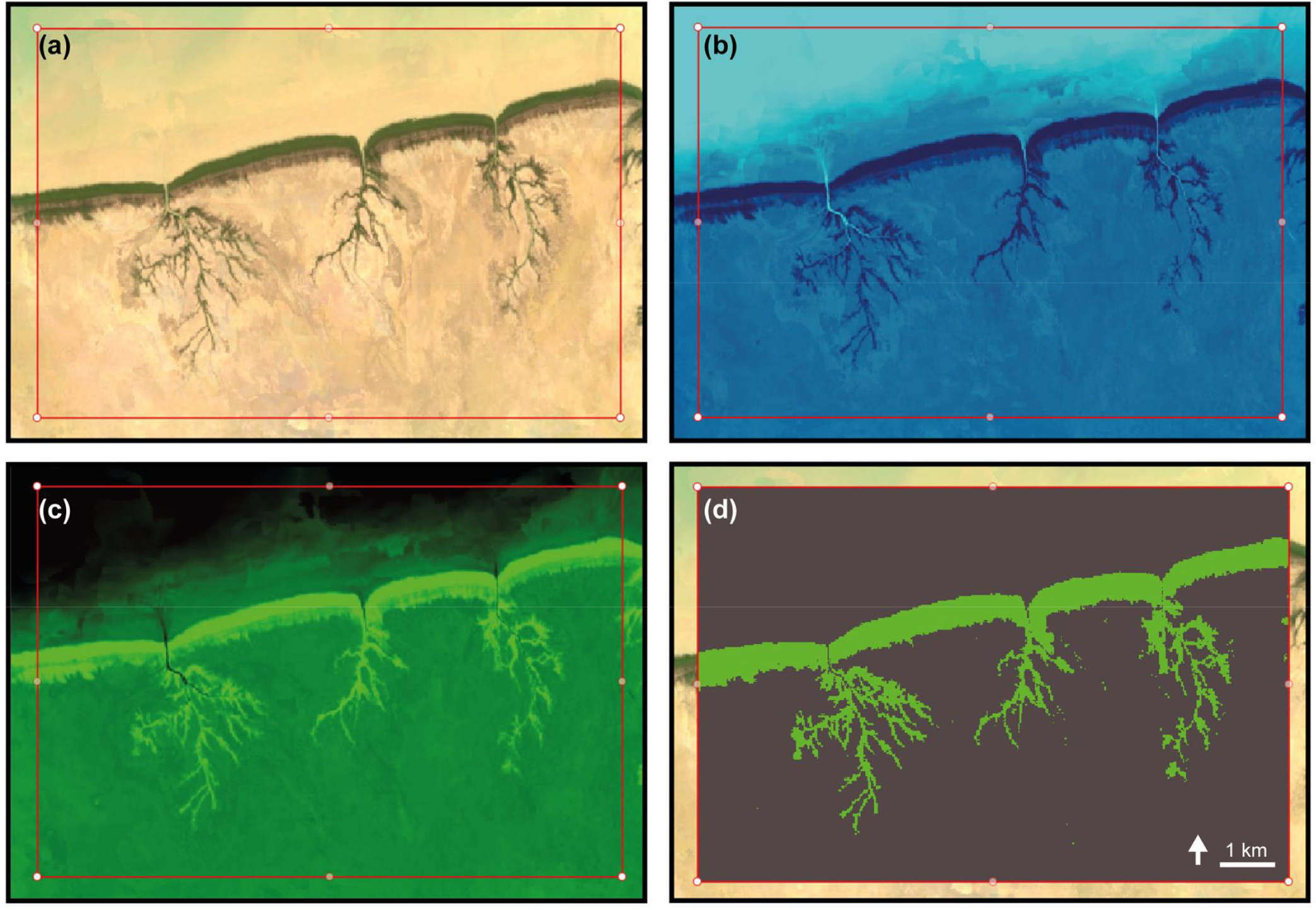
Workflow demonstrating the use of REMAP to map of a single ecosystem type, mangroves of the Gulf of Carpentaria, Australia. The panels show (a) the Landsat 8 OLI 3-year composite base layer from which all Landsat indices available in REMAP are calculated, (b) the Normalized Differenced Water Index (NDWI), (c) the Normalized Differenced Vegetation Index (NDVI), and (d) the final classified map of the distribution of mangroves in the region of interest (red box).

**Figure 3.**
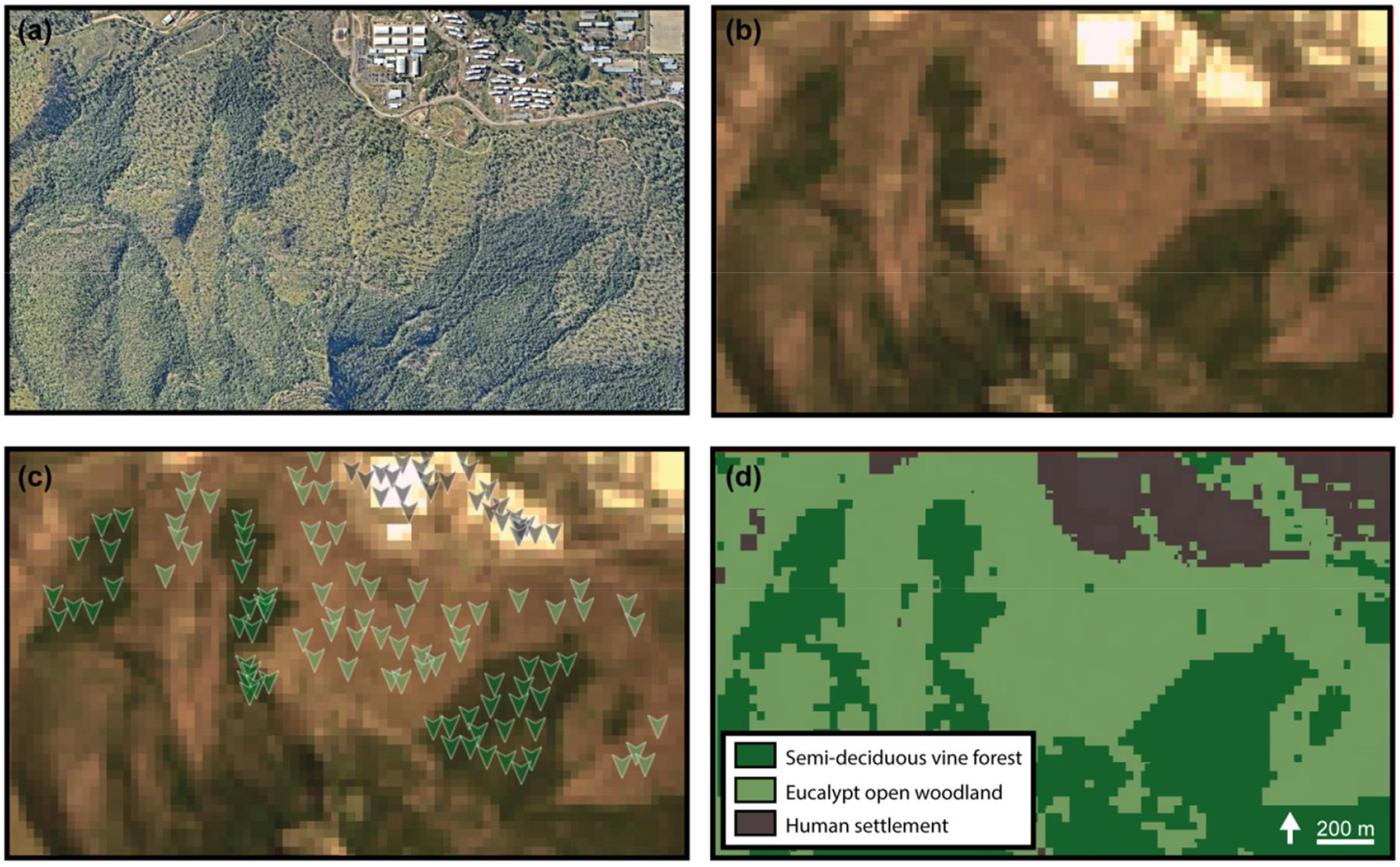
Demonstration of the use of Remap to classify ecosystem types, Mount Stuart, Queensland, Australia. (a) High resolution aerial photograph, (b) the 2017 Landsat OLI image composite, (c) training data used to produce the final 3-class map, and (d) the final classified map of the distribution of ecosystems in the focal region. Aerial photography in panel (a) copyright 2017 Nearmap Australia Pty Ltd.

**Figure 4.**
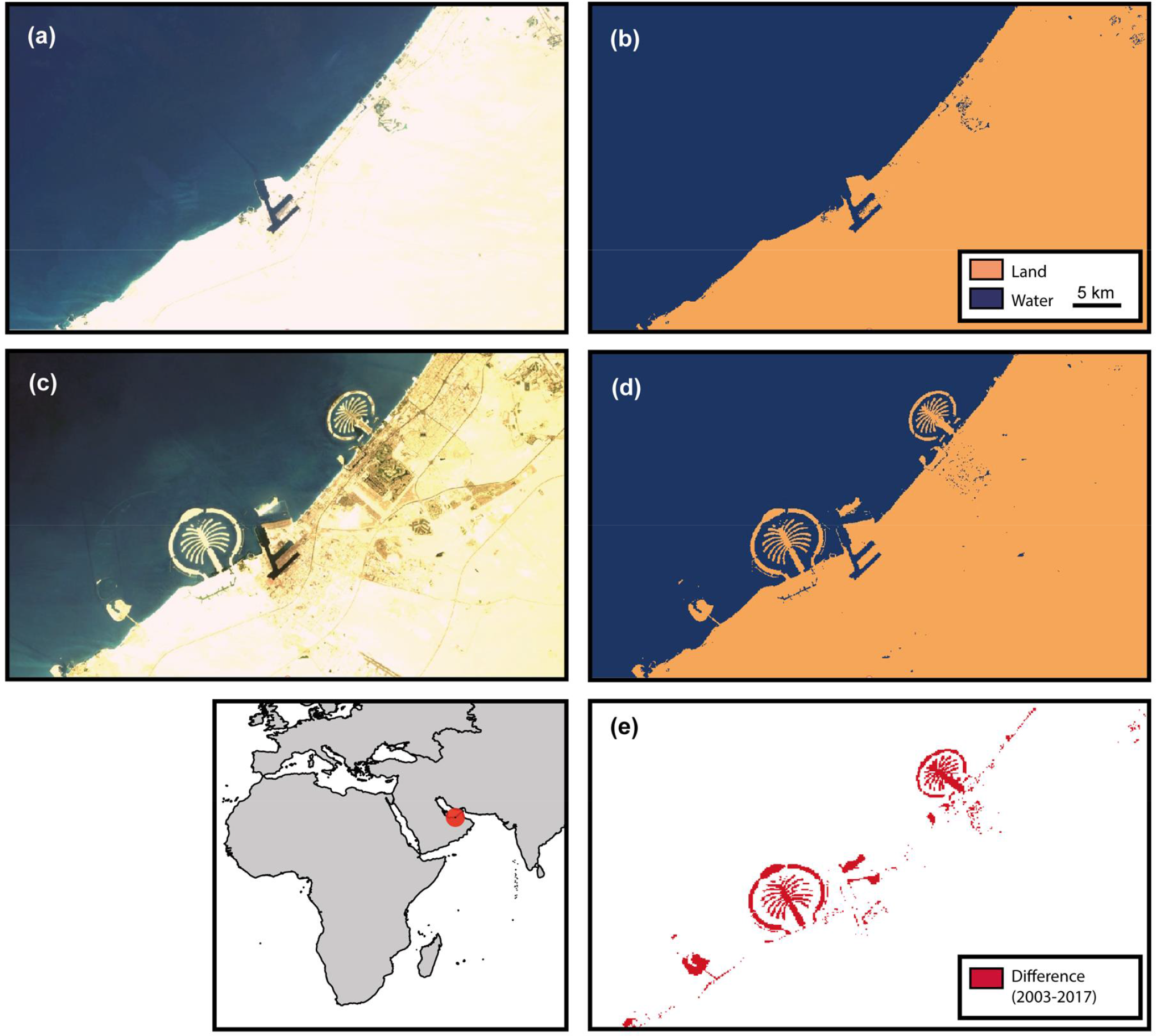
The use of REMAP to identify cover change between 2003 and 2017, Dubai. The classified land-water maps developed from (a) the 2003 global Landsat mosaic and (b) 2003 land-water classification (c) 2017 global Landsat mosaic and (d) 2017 land-water classification. (e) image differencing allows areas of coastal reclamation to be mapped and quantified. Refer to Figure 1 and Appendix A for a deforestation example.

## DISCUSSION

REMAP is a fast, user-friendly approach to developing land cover maps from freely available remote sensing data and its outcomes can be accepted if the accuracies of classifications meet the expectations of the users. Our case studies indicate that such accuracies can be achieved in REMAP but these depend upon the accuracy of the training data. By utilizing the geospatial storage and analysis capacity of the GEE, REMAP allows users with no prior knowledge in remote sensing and analysis to develop maps directly within a web-browser. This enables mapping to be undertaken in regions by locally-responsible individuals and organisations where computing infrastructure is scarce or the quality of internet connections do not allow the download of remote sensing data for local analyses. Indeed, REMAP is particularly useful for participatory mapping projects, expert elicitation and engagement with a wide-range of environmental stakeholders.

We acknowledge that REMAP has several limitations. Most notably, the ability of REMAP to produce accurate maps is limited by the quality of the training data, the accuracy of the predictors, and the suitability of the predictor set for distinguishing land cover classes and. Further development of the REMAP application will therefore include incorporating a greater number of relevant predictor data layers, such as climate maxima and minima. Future work will also focus on (i) incorporating new global image composites from the same or different years to allow monitoring of land use and cover change with higher temporal resolution or selection of specific time frames by users, (ii) utilizing all relevant and available satellite imagery (e.g. Sentinel 2), (iv) improving the user experience through the provision of more analysis tools (e.g. image differencing), and (v) improving the application for use in collecting field data and producing maps in mobile devices.

In conclusion, we have developed REMAP to make remote sensing accessible to a very wide audience with the aim of broadening the use of classified maps in ecosystem monitoring and conservation programs, and to help support the conservation of natural environments. We expect REMAP to extend the ability of volunteers, students, scientists and managers to assess the extent of land cover changes and implement conservation actions to reduce the loss of natural ecosystems.

## ACKNOWLEDGEMENTS

The project was supported by a Google Earth Engine Research Award and an Australian Research Council Linkage Grant LP130100435, co-funded by the International Union for the Conservation of Nature, MAVA Foundation, NSW Office of Environment and Heritage, and the South Department of Environment, Water and Natural Resources. We particularly wish to thank David Thau for advice throughout the project and the Google Earth Engine team for developing the Google Earth Engine (https://earthengine.google.com), without which this application would not be possible.

## AUTHOR CONTRIBUTIONS

N.J.M, D.A.K and R.M.L conceived the project. N.J.M and J.H.W. developed the remote sensing classification approach. J.H.W., D.S. and N.J.M wrote the application code and website. N.J.M wrote the manuscript with input from all coauthors.

## DATA ACCESSIBILITY

REMAP: the remote ecosystem monitoring and assessment pipeline is a freely accessible and open-source web application available at: www.remap-app.org. Data used to develop the figures in this manuscript are available at figshare (https://figshare.com/s/7125654aded6d9235f08). A snapshot of the remap source code at the time of publication is available (www.github.com/REMAPApp/REMAP). Nearmap aerial imagery courtesy of Nearmap Pty. Ltd. (© 2017 Nearmap Australia Pty. Ltd.).

## SUPPLEMENTARY MATERIAL

## Appendix A: Supplementary Data

Data used to produce Figure 1 and Figure S3

- training points (remap_points_roraimaForest.csv)
- remap workspace (remap_training_roraimaForest.json)

Data used to produce Figure 2

- training points (remap_points_carpentariaMangroves.csv)
- remap workspace (remap_training_carpentariaMangroves.json)

Data used to produce Figure 3

- training points (remap_points_mtStuart.csv)
- remap workspace (remap_training_mtStuart.json)

Data used to produce Figure 4

- training points (remap_points_Dubai_2003.csv)
- remap workspace (remap_training_Dubai_2003.json)
- training points (remap_points_Dubai_2017.csv)
- remap workspace (remap_training_Dubai_2017.json)

Data used to produce Figure S2

- training points (remap_points_chedubaMyanmar.csv)
- remap workspace (remap_training_chedubaMyanmar.json)

## Appendix B: Land cover example 2

**Figure S1.**
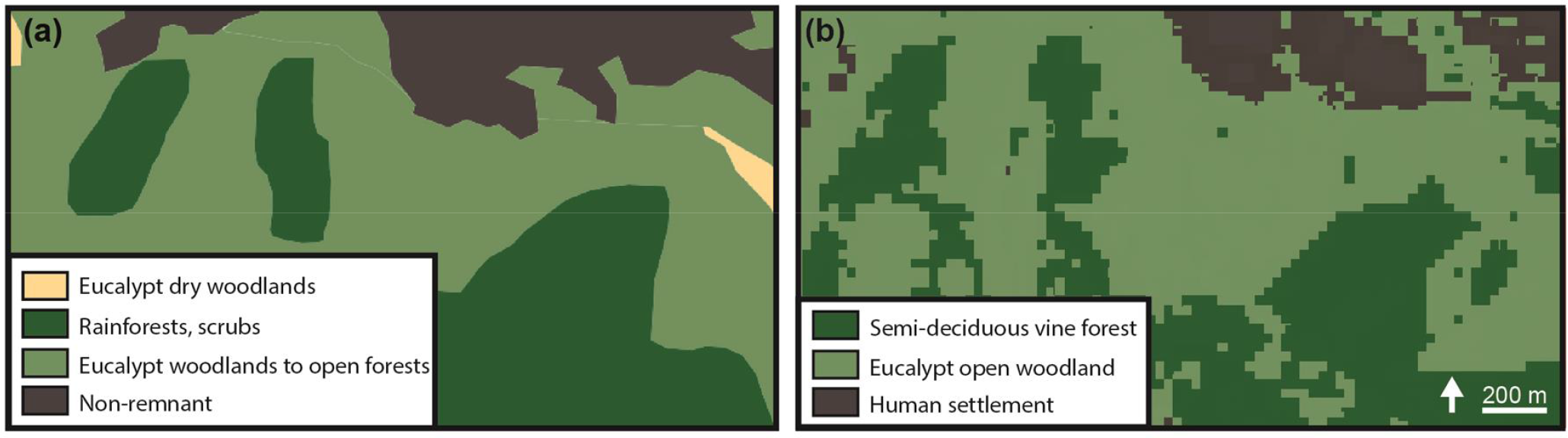
Comparison of land cover map produced by the Queensland State Government with the REMAP map shown in Figure 3, Mount Stuart, Queensland, Australia. (a) Queensland government regional ecosystem map produced from aerial photography and satellite image interpretation (Neldner *et al.* 2017; Queensland Department of Natural Resources and Mines 2017), (b) the classified map of the distribution of major ecosystems in the focal region produced with REMAP.

**Figure S2.**
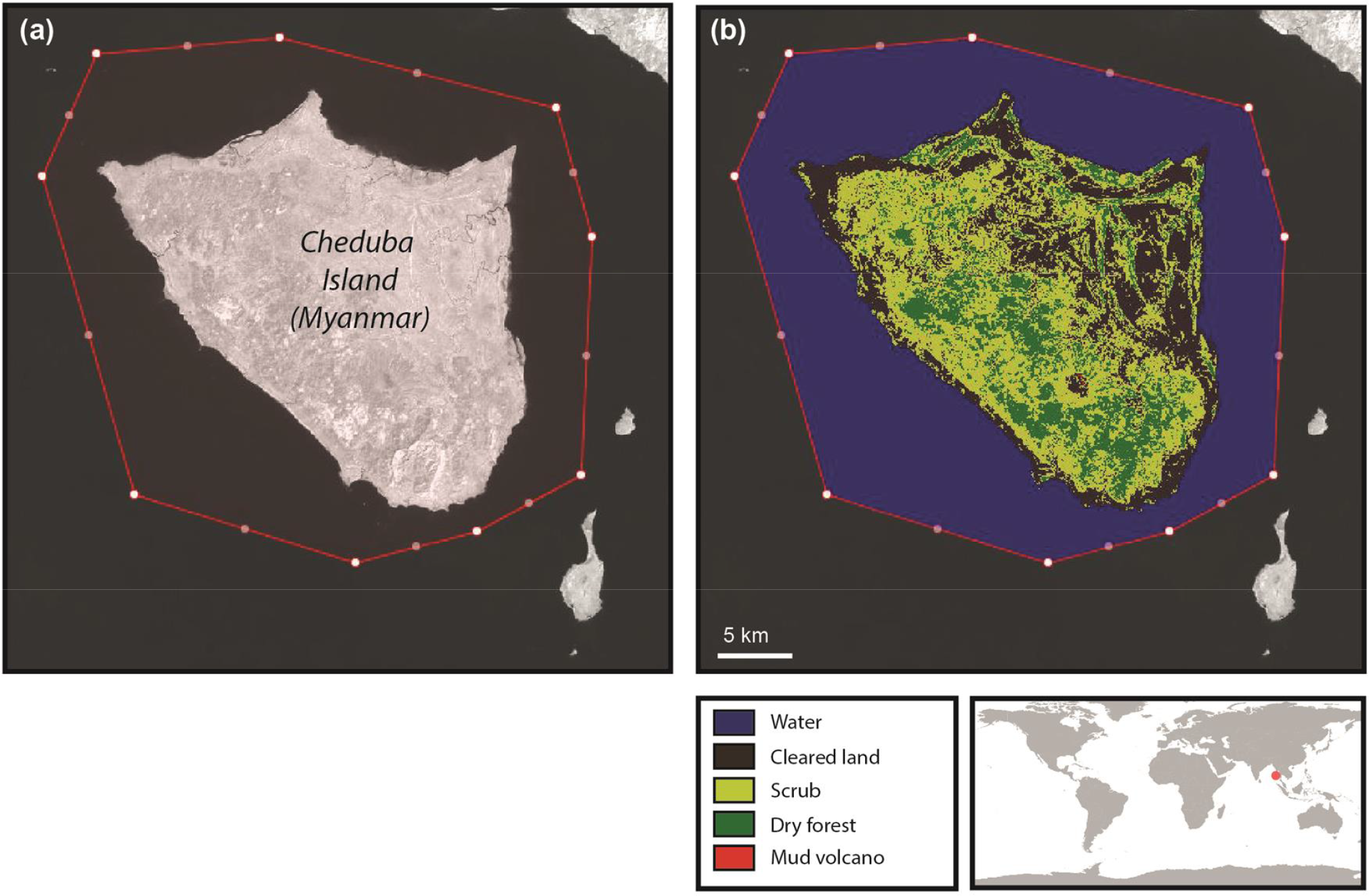
Demonstration of the use of REMAP to classify land cover types in Cheduba Island, Myanmar. The focal region for which the classification is implemented is shown by the red polygon.

**Figure S3.**
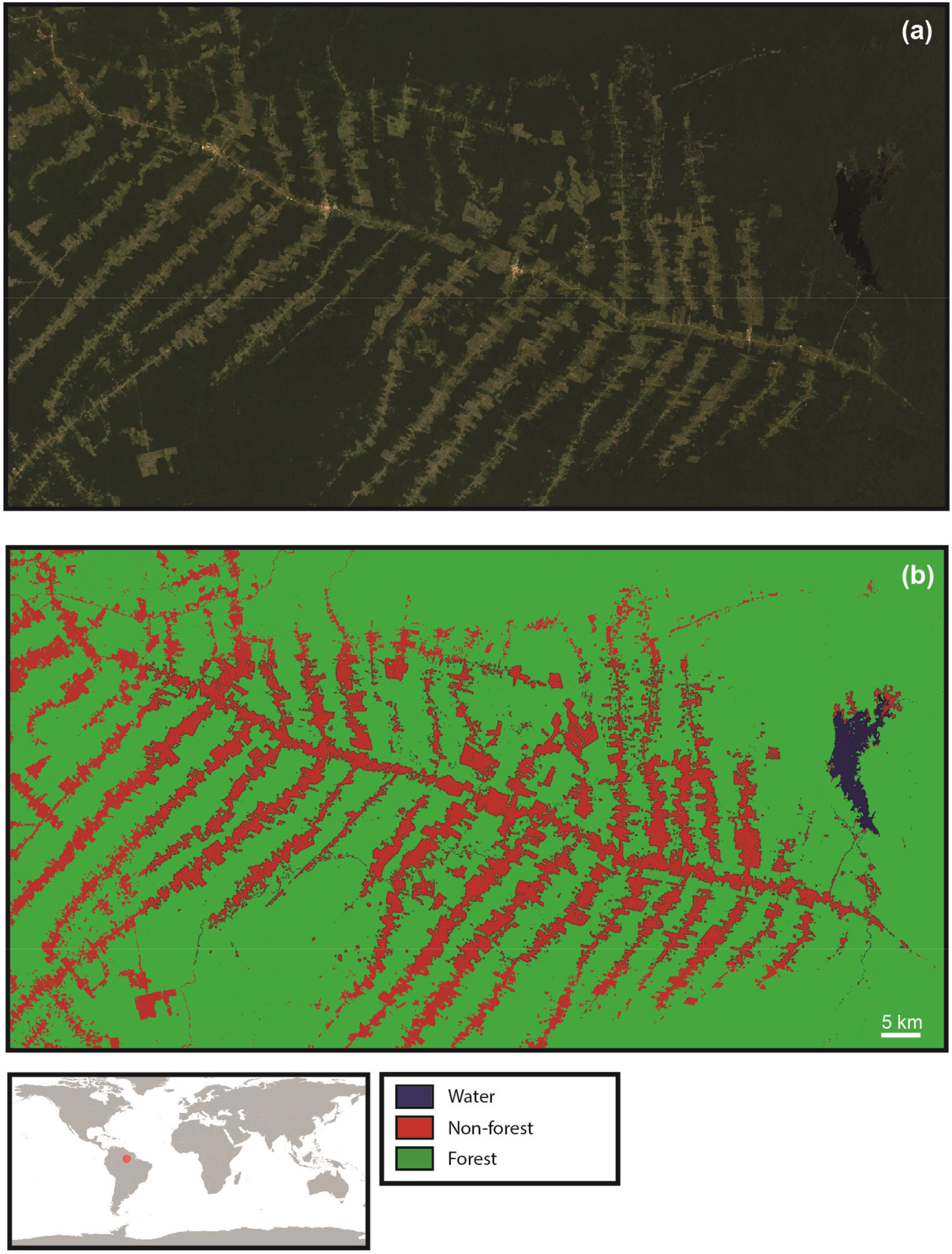
Demonstration of the use of REMAP to map deforestation in the Roraima area of Brazil.

